# Prevalent fast evolution of genes involved in heterochromatin functions

**DOI:** 10.1101/2024.03.03.583199

**Authors:** Leila Lin, Yuheng Huang, Jennifer McIntyre, Ching-Ho Chang, Serafin Colmenares, Yuh Chwen G. Lee

## Abstract

Heterochromatin is a gene-poor and repeat-rich genomic compartment universally found in eukaryotes. Despite its low transcriptional activity, heterochromatin plays important roles in maintaining genome stability, organizing chromosomes, and suppressing transposable elements (TEs). Given the importance of these functions, it is expected that the genes involved in heterochromatin regulation would be highly conserved. Yet, a handful of these genes were found to evolve rapidly. To investigate whether these previous findings are anecdotal or general to genes modulating heterochromatin, we compile an exhaustive list of 106 candidate genes involved in heterochromatin functions and investigate their evolution over short and long evolutionary time scales in *Drosophila.* Our analyses find that these genes exhibit significantly more frequent evolutionary changes, both in the forms of amino acid substitutions and gene copy number change, when compared to genes involved in Polycomb-based repressive chromatin. While positive selection drives amino acid changes within both structured domains with diverse functions and intrinsically disordered regions (IDRs), purifying selection may have maintained the proportions of IDRs of these proteins. Together with the observed negative associations between evolutionary rates of these genes and genomic TE abundance, we propose an evolutionary model where the fast evolution of genes involved in heterochromatin functions is an inevitable outcome of the unique functional roles of heterochromatin, while the rapid evolution of TEs may be an effect rather than cause. Our study provides an important global view of the evolution of genes involved in this critical cellular domain and provides insights into the factors driving the distinctive evolution of heterochromatin.

## Introduction

Heterochromatin, first discovered as the darkly stained chromosomal regions that remain condensed throughout the cell cycle (Heitz 1928), is a distinct cytological domain that is conserved across eukaryotic cells (Liu et al. 2020). While a small fraction of heterochromatin was found to be cell-type specific, or “facultative”, the majority of these chromosomal blocks remain condensed across different cell types and are known as “constitutive” heterochromatin (Heitz 1928). Constitutive heterochromatin (referred to as “heterochromatin” for simplicity hereafter) is usually located around centromeres and telomeres (Janssen et al. 2018), and its underlying DNA is depleted of functional genes. Instead, heterochromatin is mainly composed of repetitive sequences, including satellite repeats (Peacock et al. 1978) and transposable elements (TEs) (Hoskins et al. 2007; Hoskins et al. 2015). Accordingly, heterochromatin is oftentimes assumed to be functionally inert and nicknamed the “dark matter” of the genome. Yet, studies have found heterochromatin playing critical roles in many chromosomal functions, such as maintaining genome stability, mediating chromosome segregation, and ensuring proper DNA repair of repeat-rich sequences, across eukaryotes (reviewed in (Feng and Michaels 2015; Allshire and Madhani 2018; Janssen et al. 2018; Kendek et al. 2021)). Not surprisingly, disruption of heterochromatin functions has been linked to various diseases (Hahn et al. 2010), aging progression (Villeponteau 1997; J.-H. Lee et al. 2020), and cancer (Janssen et al. 2018).

The critical functions and inter-species conservation of heterochromatin domains would naturally lead to the expectation that genes modulating the function of heterochromatin should be highly conserved. Surprisingly, however, a handful of these genes show non-neutral evolution of their amino acid sequence and rapid turnover of gene copy number, both of which are consistent with a history of positive selection. The evolution of some of these genes was of interest due to them being core structural components of heterochromatin, such as the Heterochromatin Protein (HP1) family (Levine et al. 2012; Helleu and Levine 2018) and its paralogs (Vermaak et al. 2005; Ross et al. 2013). The rest of the genes were observed to have roles in the evolution of small RNA pathways (Obbard et al. 2006; Obbard et al. 2009), transcription factors (Kasinathan et al. 2020), or germline stem cells (Flores et al. 2015), as well as speciation (Barbash et al. 2003). Because satellite repeats (Henikoff et al. 2001) and TEs (Cosby et al. 2019) are enriched in heterochromatic sequence and can defy Mendel’s law to increase their transmission to the next generation, an arms race between heterochromatic sequence and host genes suppressing such selfish behaviors has been a common theme explaining the rapid evolution of these genes (e.g., (Vermaak et al. 2005; Satyaki et al. 2014; Helleu and Levine 2018; Brand and Levine 2021)). Further supporting such conjecture, the composition, abundance, and location of repetitive sequences in heterochromatin are found to diverge rapidly even between closely related species (Wei et al. 2014; Wei et al. 2018; Kim et al. 2021; de Lima and Ruiz-Ruano 2022).

However, this handful of genes represents only a small fraction of genes involved in modulating heterochromatin functions. The unique and essential functions of heterochromatin depend on its specific enrichment of repressive di– and tri-methylation at histone H3 Lysine 9 (H3K9me2/3, (Allshire and Madhani 2018)) and the “reader” of these marks, HP1a, which serves as the foundational structural component of heterochromatin (Eissenberg and Elgin 2014) and leads to DNA compaction (Verschure et al. 2005) and transcriptional suppression (Li et al. 2003). Interestingly, the binding of HP1a to H3K9me2/3 further recruits “writers” of H3K9me (histone methyltransferase, (Schotta 2002)), and this positive feedback mechanism between readers and writers leads to the propagation of such repressive epigenetic marks independent of the underlying DNA sequence (Allshire and Madhani 2018). This unique ability of heterochromatin to “spread” is best exemplified by the Position Effect Variegation (PEV) first discovered in *Drosophila* (Muller 1930)—the mosaic expression of euchromatic genes translocated to heterochromatin-proximal regions due to the stochastic spreading of repressive H3K9me2/3 from heterochromatin (Girton and Johansen 2008; Elgin and Reuter 2013). By studying mutants that either enhance or weaken PEV, many genes involved in heterochromatin functions were identified. These include not only the structural (e.g., (Shaffer et al. 2006)) and enzymatic (e.g., (Czermin et al. 2001)) components for heterochromatin, but also those that antagonize the initiation and/or maintenance of heterochromatin (e.g., writers of antagonizing histone modifications; (Bao et al. 2007)). Still, other genes were discovered through their co-localization with heterochromatin domain either cytologically (e.g., (Swenson et al. 2016)) or epigenomically (e.g., (Alekseyenko et al. 2014)) and were subsequently identified to be involved in heterochromatin function.

Given the large numbers and vastly diverse functional roles of genes involved in heterochromatin function, we conducted an expansive survey to determine whether the previously reported rapid evolution of a small subset of these genes is anecdotal or a common feature of proteins involved in the functions of this unique chromatin environment. To do so, we compiled an exhaustive list of candidate genes that have been shown, or are likely, to be involved in heterochromatin functions, including PEV modifiers, histone-modifying enzymes influencing H3K9me2/3 enrichment, and genes whose protein products localize to heterochromatin. Our investigation finds that these genes evolve exceptionally fast, a pattern that is not general to genes interacting with other repressive chromatin marks and suggests selective pressure unique to constitutive heterochromatin. We further dissect the domains and protein properties targeted by positive selection and specifically test the premise that the rapid evolution of genes involved in heterochromatin functions could be driven by fast-changing repetitive sequences. Based on our findings, we propose an evolutionary model where the rapid changes in these genes are the unavoidable consequence of the unique functional roles of heterochromatin, and that the evolution of repetitive sequences may be the consequence, instead of the cause. Our study provides an important global view of the evolution of genes involved in this critical cellular domain and sheds light on the drivers behind the unique evolution of heterochromatin.

## Results

### Identification of candidate genes involved in heterochromatin functions

With the goal of identifying the global evolutionary patterns for genes involved in the functions of heterochromatin, we selected candidate genes based on three criteria to maximize inclusivity (**Table S1**). The first category contains genes demonstrated to modulate PEV (Girton and Johansen 2008; Elgin and Reuter 2013; Swenson et al. 2016). Genes with mutations weaken PEV and, therefore, heterochromatin functions are known as Suppressors of Variegation (*Su(var)*), while those that augment PEV are Enhancers of Variegation (*E(var)*). Here, we included 59 and seven previously identified *Su(var)s* and *E(var)s,* respectively.

The second category focuses on histone-modifying enzymes that can influence the enrichment levels of H3K9me2/3. Enrichment of H3K9me2/3 requires “eraser” proteins that first remove antagonizing active marks, such as H3K9ac (deacetylases, (Czermin et al. 2001)) and H3K4me3 (demethylase, (Rudolph et al. 2007)). This is followed by writer proteins depositing H3K9me2/3 (H3K9 methyltransferases, (Schotta 2002)). We consider these histone-modifying enzymes to enhance H3K9me2/3 enrichment and, thus, heterochromatin functions. On the contrary, erasers for H3K9me2/3 (H3K9 demethylase, (Herz et al. 2014)), as well as writers for acetylation (acetyltransferase, (Kuo and Andrews 2013)) and S10 kinase (Deng et al. 2008), are known to antagonize the deposition and maintenance of H3K9me2/3. H4K20me3 is another conserved hallmark of heterochromatin (Schotta et al. 2004), and we included the corresponding methyltransferases. In total, our list contains 16 and seven histone-modifying enzymes that enhance or weaken heterochromatin functions, respectively.

The last category includes genes whose protein products co-localize with heterochromatin, suggesting they play roles in this subnuclear compartment. We surveyed the literature for cytology-based evidence of co-localization with HP1a through either immunofluorescence (e.g., (Greil et al. 2007)) or live imaging (e.g., (Swenson et al. 2016)) as well as epigenomics through Chromatin Immunoprecipitation (ChIP, e.g., (Alekseyenko et al. 2014; Kasinathan et al. 2020)). Because a previous comprehensive survey found that some HP1a-binding proteins identified through Immunoprecipitation-Mass Spectrometry (IP-Mass Spec) are not necessarily enriched in heterochromatin cytologically (Swenson et al. 2016), genes implicated in HP1a-binding but lacking localization evidence are not included in our analysis. Hereafter, we referred to this largest category with 63 genes as “heterochromatin proteins.”

In total, we studied the evolution of 106 genes involved in heterochromatin functions, which we termed “heterochromatin-related genes” hereafter (see **Figure 1A** for the number of genes in each category and **Table S1** for a full list of genes and references). It is worth noting that 43 candidate genes belong to more than one category because these three categorizations are not mutually exclusive (**Table S1**). We consider genes in categories of *Su(var)s*, histone modifying enzymes enhancing H3K9me2/3 or H4K20me3 enrichment, or heterochromatin proteins as “mediating” heterochromatin functions, while those defined as *E(var)s* or histone modifying enzymes weakening H3K9me2/3 enrichment as “antagonizing” heterochromatin functions. To determine whether the evolutionary histories of heterochromatin-related genes are exceptional, we compared them with Polycomb group genes (Kassis et al. 2017), whose protein products are enriched at facultative heterochromatin. Even though also generally associated with transcriptional suppression, facultative heterochromatin is restricted to developmentally regulated genes in euchromatin and is enriched for another type of repressive histone modification (H3K27me3) (Bannister and Kouzarides 2011; Bell et al. 2023). Polycomb genes thus allowed us to determine whether the exceptional evolutionary histories, if any, of heterochromatin-related genes are due to their involvement in a repressive chromatin environment or specifically for heterochromatin structure.

**Figure 1.**
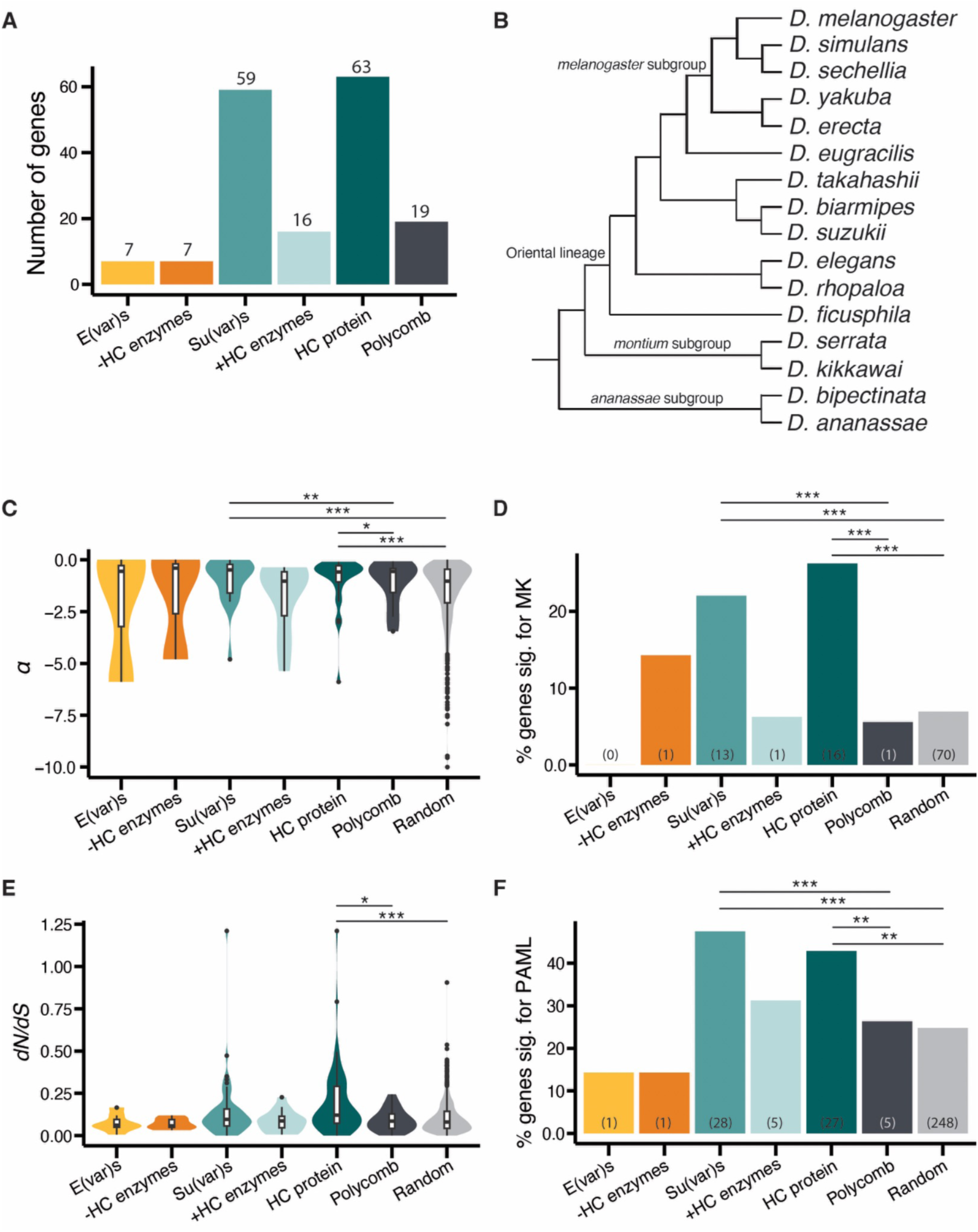
Positive selection on protein sequences of heterochromatin-related genes found by evolutionary tests at both short– and long-evolutionary time scales. (A) Barplot showing the number of genes in each category of heterochromatin-related genes, the Polycomb control genes, and a set of one thousand random genes. Those expected to mediate or antagonize heterochromatin functions are in hues of green and orange, respectively. –HC enzyme: histone-modifying enzymes weakening H3K9me2/3 enrichment; +HC enzyme: histone-modifying enzymes enhancing H3K9me2/3 enrichment. (B) Phylogenetic tree for the species included in the study. Note that the branch lengths are not to scale. (C and E) Violin plots for α, the proportion of amino acid substitutions fixed by positive selection inferred by the MK test (C) and *dN/dS* ratio, the relative rates of nonsynonymous substitutions (E), for different categories of genes. Numbers of significant genes are in parentheses. (D and F) Barplots showing the proportion of genes with significant MK tests (D) and accelerated rates of *dN/dS* driven by positive selection (F). The number of genes in each category is either on top of the barplot (A) or in parenthesis (D and F). *Mann-Whitney Test* (C and E) and *Binomial test* (D and F): \**p* < 0.05, \*\**p* < 0.01, \*\*\**p* < 0.001

### Heterochromatin-related genes experience pervasive positive selection on protein sequences across short and long-time scales

We performed evolutionary genetic tests to identify heterochromatin-related genes with signatures of positive selection at two time scales—contrasting the polymorphism within *D. melanogaster* and divergence between *D. melanogaster* and *D. simulans* (∼4 million years divergence between lineages) under the framework of unpolarized McDonald-Kreitman tests (MK tests, (McDonald and Kreitman 1991)), and phylogenetic analysis by maximum likelihood (PAML) tests (Yang 2007) using 16 *Drosophila* species with an estimated divergence time of ∼25 million years (Suvorov et al. 2022) (see **Figure 1B** for studied species). In addition to comparing the evolution of heterochromatin-related genes to that of polycomb genes, we made the contrast to the evolution of a thousand randomly sampled genes to represent genome-wide estimates (referred to as “random genes” hereafter).

On a short evolutionary time scale, we first estimated the proportion of amino acid substitutions that might have been driven by positive selection (α, (Smith and Eyre-Walker 2002)). Heterochromatin-related genes, as a group, have significantly larger α than those of Polycomb (*Mann-Whitney U tests, p =* 0.0244*, median =* 0.0847 (heterochromatin-related genes) and –0.424 (Polycomb)) and random genes (*Mann-Whitney U tests, p* < 10^-8^, *median* = – 0.516 (random genes)). Breaking down our candidate genes according to functions revealed that most categories have larger α than Polycomb control or random genes, even though the comparisons are statistically significant only for *Su(var*) and heterochromatin protein (**Figure 1C**). These observations suggest the possibility that heterochromatin-related genes experienced frequent positive selection. Consistently, we identified that 21 out of 104 (20.2%) heterochromatin-related genes with sufficient polymorphism data to perform MK tests have evidence of adaptive evolution (rejection of the null hypothesis and an excess of nonsynonymous fixed differences between species). This is a significantly larger proportion than that of the Polycomb genes (5.56%; *Fisher’s Exact Test*, *p =* 0.191; *Binomial test*, *p* < 10^-6^, odds ratio = 4.30) and random genes(6.94%, *Fisher’s Exact Test*, *p <* 10^-3^; *Binomial test*, *p <* 10^-5^, odds ratio = 2.90), while we found no difference in this proportion between Polycomb genes and random genes (*Fisher’s Exact Test* and *Binomial test*, *p =* 1). Specifically, *Su(var)* (22.0%) and heterochromatin proteins (26.2%), both of which are expected to mediate heterochromatin functions, have much larger proportions of genes with evidence of positive selection than either Polycomb or random genes (**Figure 1D**). Indeed, candidate genes mediating heterochromatin functions, as a group, show significantly more evidence of adaptive evolution than the Polycomb (20.2%, *Fisher’s Exact Test*, *p* = 0.190; *Binomial test*, *p* < 10^-6^, odds ratio = 4.30) as well as random genes (*Fisher’s Exact Test*, *p* < 10^-3^; *Binomial test*, *p* < 10^-4^, odds ratio = 2.91). It is worth noting that the number of codons involved in the MK tests is similar across categories of genes (*Mann-Whitney U tests, p >* 0.05 for all comparisons), suggesting that these observations are unlikely driven by differences in the statistical power of the MK tests.

On a long evolutionary time scale, we estimated the relative rates of amino acid substitution (nonsynonymous divergence/synonymous divergence, *dN/dS*) across 16 studied *Drosophila* species. There is an overall trend that heterochromatin-related genes have larger *dN/dS* ratios than the Polycomb control genes (medians = 0.0974 (heterochromatin-related genes) *v.s.* 0.0852 (Polycomb controls)) and random genes (medians = 0.0802), and this difference is statistically significant for comparisons to random genes (*Mann-Whitney U test, p* = 0.167 (compared to Polycomb control) and 0.0032 (compared to random genes). Nevertheless, only the category of “heterochromatin protein,” the largest category, has a statistically larger *dN/dS* ratio than that of the Polycomb genes (*Mann-Whitney U test, p* = 0.0204) and random genes (*Mann-Whitney U test, p* < 10^-5,^ **Figure 1E**). To test whether the changes in *dN/dS* ratios of heterochromatin-related genes over the phylogenetic tree might have been driven by positive selection, we compared the log-likelihood for two alternative models estimating *dN/dS* at each site: the null model that assumes *dN/dS* ratio is beta-distributed and not greater than one (M8a) and the alternative model in which *dN/dS* ratio is allowed to be greater than one (M8 models; (Yang 2007)). A significantly larger likelihood of the alternative model is consistent with a history of positive selection acting on the gene (Swanson et al. 2003). We found that the substitution patterns of 42 heterochromatin-related genes (39.6%) better fit the alternative model, suggesting frequent positive selection acting on them. This proportion is significantly larger than that of Polycomb genes (26.3%; *Fisher’s Exact Test*, *p =* 0.314; *Binomial test*, *p =* 0.00276, odds ratio = 1.84) and the random genes (24.8%; *Fisher’s Exact Test*, *p =* 0.00159; *Binomial test*, *p =* 0.000975, odds ratio = 1.99), while there was no difference in this proportion between the Polycomb and random genes (*Fisher’s Exact Test*, *p =* 0.795; *Binomial test*, *p* = 0.796).

Similar to observations made on a short evolutionary time scale, *Su(var)s* and heterochromatin proteins have stronger evidence of positive selection compared to Polycomb and random genes (**Figure 1F**), as do all genes mediating heterochromatin functions (41.4%, *Fisher’s Exact Test*, *p* = 0.305; *Binomial test*, *p* = 0.00125, odds ratio = 1.97 (compared to Polycomb control) and *Fisher’s Exact Test*, *p* = 0.0115; *Binomial test*, *p* = 0.000272, odds ratio = 1.67 (compared to random genes)). Interestingly, while histone-modifying enzymes that enhance heterochromatin, as a group, do not exhibit exceptional rates of protein evolution, likely due to the low statistical power associated with its small number of genes, *all* methyltransferases for H3K9me2/3 (*Su(var)3-9, egg, G9a*) show accelerated rates of protein evolution (**Table S1**), which provides clues for the possible source of positive selection acting on heterochromatin-related genes (see Discussion). Overall, we found that heterochromatin-related genes, especially those mediating heterochromatin functions, experienced frequent positive selection on their protein sequence at both short– and long-time scales, even when compared to Polycomb genes. This observation suggests that the widespread adaptive evolution of heterochromatin-related genes is likely driven by selection acting on their functions specific to the heterochromatin environment, rather than by general selective pressure acting on genes interacting with repressive epigenetic marks.

### Rapid evolution of heterochromatin-related genes also manifests as changes in gene copy number

In addition to changes in amino acid sequences, the rapid evolution of genes can be in the form of changes in gene copy number (Hastings et al. 2009), and dramatic turnover of gene copy number for a small set of heterochromatin-related genes was previously reported (Levine et al. 2012; Lewis et al. 2016; Lee et al. 2017; Helleu and Levine 2018). Changes in gene copy number may hold particular significance for heterochromatin functions—several enzymatic and structural proteins of heterochromatin have been shown to exhibit dosage-dependent effects (Elgin and Reuter 2013). Accordingly, changes in gene copy number of heterochromatin-related genes could have immediate functional consequences.

We performed reciprocal *BLAST* searches using *D. melanogaster* amino acid sequences as queries to identify homologs and paralogs in other species, followed by manual curations (see Materials and Methods). With these, we found 18 heterochromatin-related genes having differences in gene copy number among 16 *Drosophila* species studied (17.0% of the heterochromatin-related genes), with 17 genes having gains of copies and one gene loss when compared to *D. melanogaster.* Similar to our analyses for the evolution of amino acid sequences (**Figure 1**), the proportion of heterochromatin-related genes with copy number variation (CNV) is significantly more than that of Polycomb genes (5.26%, *Fisher’s Exact Test*, *p* = 0.302, *Binomial test*, *p* < 10^-4^, odds ratio = 3.68). *Su(var)* genes and heterochromatin proteins again exhibit exceptionally large proportions of genes with CNV (**Figure 2A**). CNV of several of these genes have been studied before (e.g., *HP1* (Levine et al. 2012), *AGO2* (Lewis et al. 2016)*, Cav* and *Nap-1* (Lee et al. 2017), and *mh* (Brand et al. 2024)) and our following discussions mainly focused on those first identified by our study.

**Figure 2.**
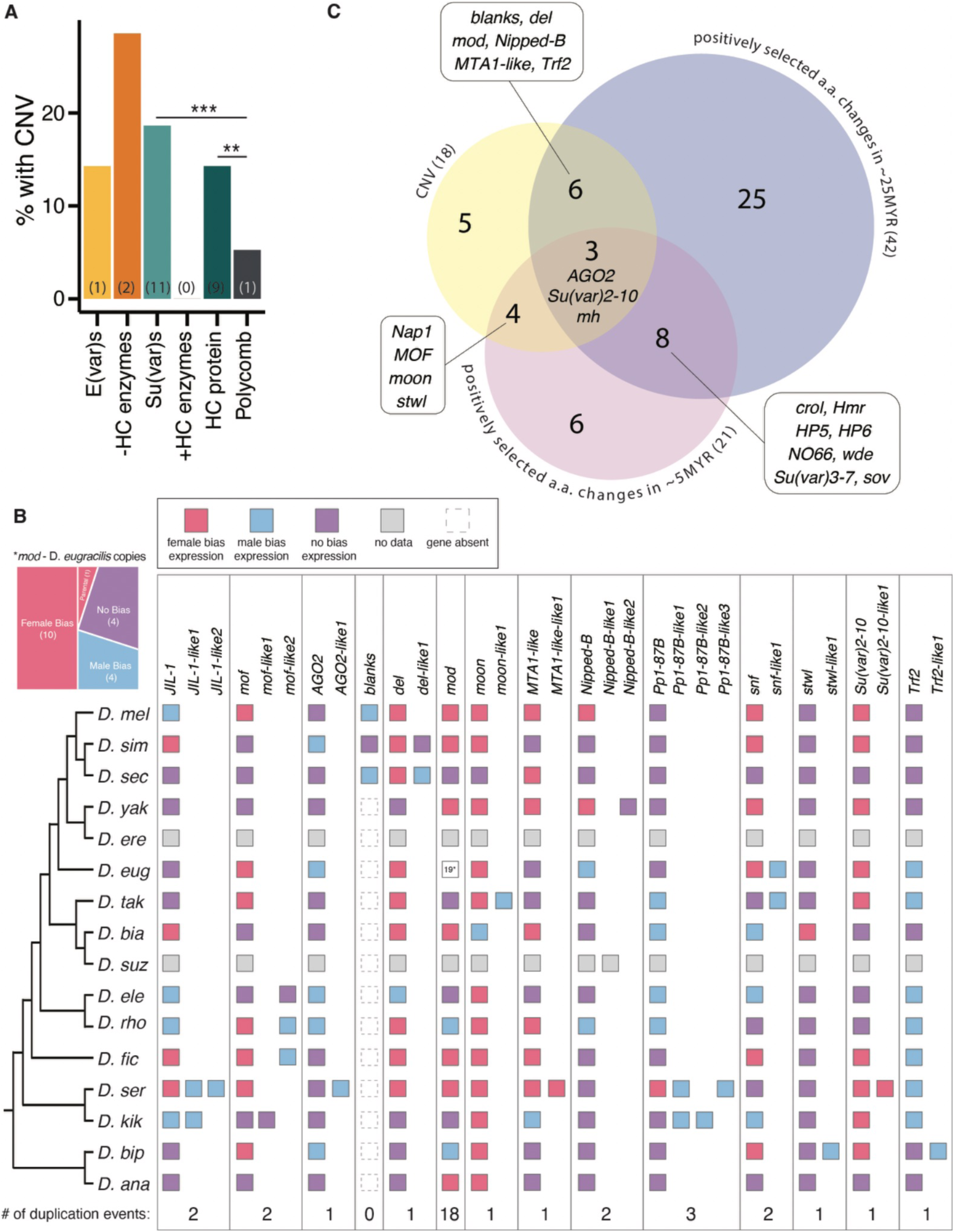
Rapid evolution of heterochromatin-related genes manifested in multiple forms. (A) Barplot showing the proportion of heterochromatin-related genes and Polycomb control genes with CNV. The numbers of genes with CNV are in parentheses. (B) Summary of the duplication and loss events as well as sex-biased expression for heterochromatin-related genes identified to have CNV. Each column represents one orthologous copy. The number of duplication events is noted at the bottom. The colors of each square represent sex-biased expression, except for the 19 copies of *mod* duplicates in *D.eugracilis,* which is shown at the top left of figure B. (C) Venn diagram depicting the number of genes identified to have exceptional evolution with one of the three evolutionary tests as well as their overlaps, with those identified by more than one test listed. a.a.: amino acid. *Binomial tests*: \*\**p* < 0.01; \*\*\**p* < 0.001.

By using synteny to infer the orthologous relationships between duplicates in different species (see Materials and Methods), we found that most observed CNVs for heterochromatin-related genes involve a single or two duplication events (**Figure 2B, Figure S1, and Table S2**). The most notable exception is the nucleolin homolog, *mod,* which has 18 duplications in *D. eugracilis* alone. *mod* plays important roles in morphogenesis (Graba et al. 1994) and spermatogenesis (Park et al. 2023) and has been shown as a dosage-dependent *Su(var)* (Garzino et al. 1992), suggesting these identified CNVs could readily alter heterochromatin functions. In addition, we observed complex duplication/loss events leading to the CNV observed in *mof* (**Figure S2**), an acetyltransferase that influences H3K9me2/3 enrichment (Feller et al. 2015) and is known to be involved in dosage compensation (Hilfiker et al. 1997). In addition to the duplication happening on the lineage leading to *D. kikkawai,* there is likely a duplication event of *mof* in the common ancestor of the oriental lineage (see **Figure 1B**), followed by a subsequent loss in the subsets of lineages that leads to the paraphyletic presence of a *mof* duplicate (**Figure S2**).

Duplicated genes are commonly found to have male-biased expression, which is suggested as a resolution to sexual genetic conflicts (Gallach and Betrán 2011). We are interested in examining whether duplicates of heterochromatin-related genes exhibit similar trends of sex-biased expression. By using publicly available transcriptome data for the species studied, we categorized the parental ortholog and duplicated paralog (with respect to *D. melanogaster*) as male-biased, female-biased, or unbiased in expression (see Materials and Methods). With the exception of *MTA-like* and *Su(var)2-10*, most identified duplicates indeed show male-biased expression, irrespective of whether the parental copy is male or female-biased in expression (**Figure 2B**). Interestingly, for *mod* in *D. eugracilis*, the parental copy has a female-biased expression, while the 18 duplicates exhibit a mixture of male-biased, female-biased, and unbiased expression. Another notable case is *del,* which is part of a germline complex enabling the transcriptions of TE-targeting small RNAs from loci in pericentromeric heterochromatin (Mohn et al. 2014) and shows female-biased expression. While the *del* duplicate in *D. simulans* shows male-biased expression, the *orthologous* duplicated copy in the sister species *D. sechellia* has no biases in expression. In summary, the rapid evolution of heterochromatin-related genes, when compared to Polycomb genes, is also reflected in their changes in gene copy number.

### Heterochromatin-related genes with diverse functions are recurrent targets of positive selection

The three evolutionary tests we conducted detect signals of positive selection at different time scales and in different forms (amino acid sequences *v.s.* gene copy number). We are interested in investigating whether some heterochromatin-related genes may exhibit rapid evolution with multiple tests. Consistent with analyses conducted separately for different evolutionary tests, *Su(var)s* and heterochromatin proteins have an especially large proportion of genes with evidence of rapid evolution in any one of the tests (*Binomial test, p <* 0.001 for both comparisons; **SFigure 3A**), and this difference becomes even more prominent when considering genes with exceptional evolution for *more than one test* (*Binomial test, p <* 0.001 for both comparisons; **SFigure 3B**). Three genes show evidence of rapid evolution for *all* three tests (**Figure 2C**): *AGO2*, the core effector in endogenous siRNA silencing pathway that silences TEs in somatic cells (Ghildiyal et al. 2008), *Su(var)2-10*, which critically links piRNA targeting and the transcriptional silencing of TEs (Ninova et al. 2020), and *mh*, whose evolution is the focus of a recent study (Brand et al. 2024). Other heterochromatin-related genes with more than one evidence of rapid evolution (**Figure 2C**) include those related to silencing of TEs (e.g., *del, moon, sov, Trf2, wde*), dosage compensation (e.g., *Su(var)3-7*), maintenance of chromatin structure (e.g., *Nipped-B*), and structural components of heterochromatin (e.g., *HP5*). While most of these genes are either *Su(var)* or heterochromatin proteins, several histone-modifying enzymes were also found to undergo positive selection with multiple evolutionary tests, such as the H3K4me3 demethylase *NO66* and acetyltransferase *mof*. These observations suggest that positive selection recurrently acts on heterochromatin-related genes with diverse functions.

**Figure 3.**
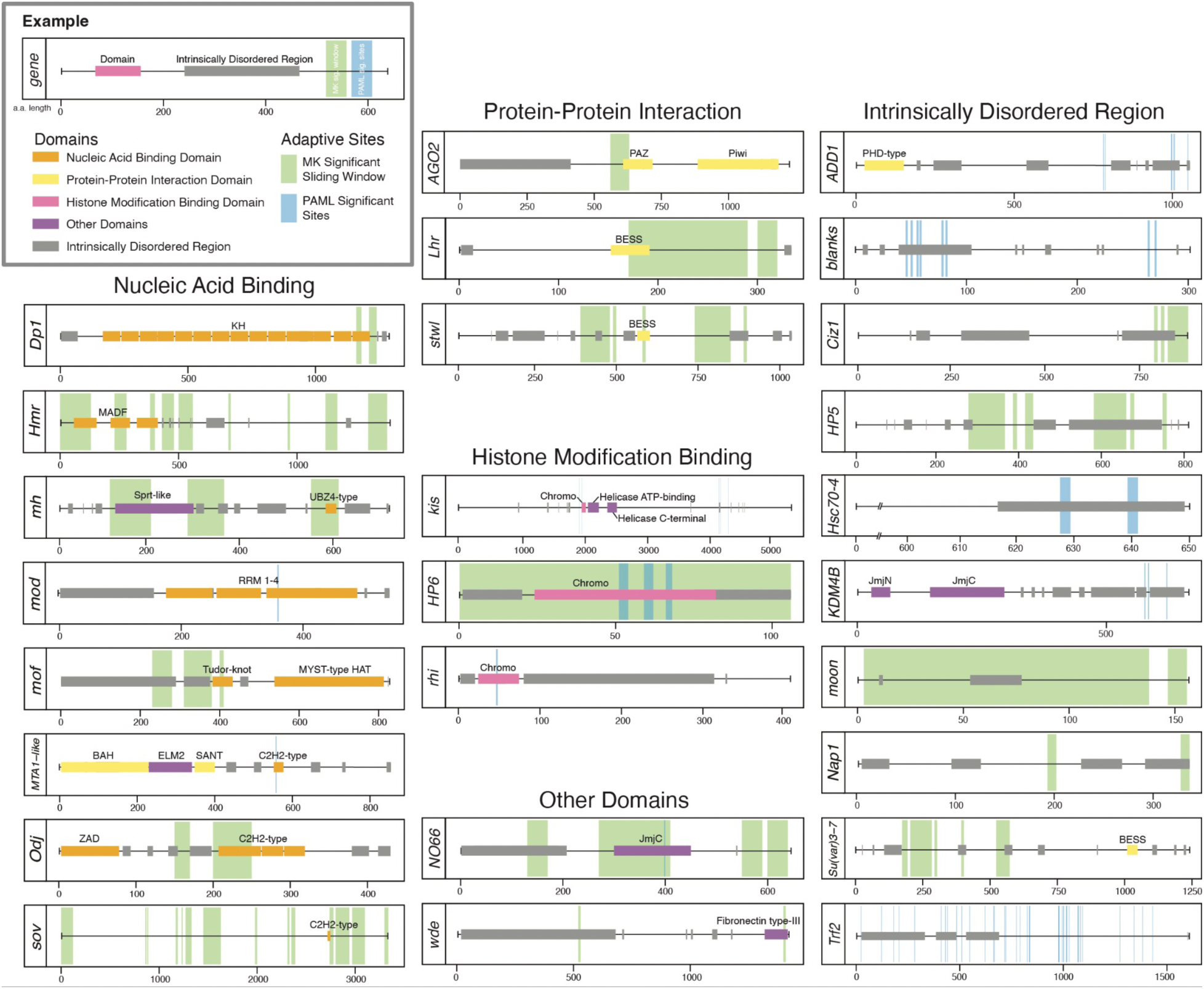
Signatures of positive selection fall within domains with diverse functions and IDRs of heterochromatin-related genes. Genes with evidence of positive selection on protein sequences are categorized according to which types of domains/regions their signatures of positive selection fall within. Annotated and/or predicted structured domains are shown as horizontal lines with the following functional categories: nucleic acid binding (orange), protein-protein interaction (yellow), histone modification binding (pink), and other domains (purple). IDRs are marked with gray. Highlighted vertical windows represent those under positive selection as identified by sliding MK tests (green) and PAML inferences (blue). Note that genes with no positively selected windows/sites overlapping with either annotated domains or IDRs were not shown.

### Positive selection targets both ordered domains and intrinsically disordered regions of proteins encoded by heterochromatin-related genes

To identify the potential evolutionary drivers for the observed rapid evolution of heterochromatin-related genes, we investigated the locations of positive selection within windows of each rapidly evolving gene. We identified windows with evidence of adaptive evolution by performing sliding MK tests and located sites with high Bayes Empirical Bayes (BEB) posterior probability of positive selection under the PAML framework (Yang et al. 2005). Locations of these windows or sites with evidence of positive selection were then contrasted with the annotated and/or predicted domains of heterochromatin-related genes (see Materials and Methods). We found that signatures of positive selection are present in domains with diverse functions (**Figure 3**), and several of them are especially pertinent to heterochromatin functions. These include chromo domains that directly interact with histone methylation (in *HP6, rhi, Kis*) and Jmjc demethylase domain in *NO66*, the H3K4me3 demethylase. We also found signatures of positive selection located within various nucleic acid binding domains (e.g., C2H2 type and UBZ4-type Zinc-Finger DNA binding domain) and domains mediating protein-protein interactions (e.g., BESS domains).

In addition to structured domains with well-characterized functions, proteins also contain regions that lack fixed three-dimensional structures, known as Intrinsically Disordered Regions (IDRs). IDRs are increasingly appreciated for playing critical roles in protein functions (Forman-Kay and Mittag 2013) and are frequently found in proteins enriched in phase-separated cellular compartments (Nott et al. 2015; Pak et al. 2016; Lin et al. 2017). Indeed, IDR-mediated phase properties have been suggested to be critical for heterochromatin functions (Larson et al. 2017; Strom et al. 2017). Accordingly, we investigated whether IDRs in heterochromatin-related genes may also have signatures of positive evolution by using flDPnn (Hu et al. 2021) to predict IDRs in *D. melanogaster* protein sequences. Interestingly, we found that signatures of rapid evolution also frequently fall within IDRs (**Figure 3**), and, for multiple rapidly evolving heterochromatin-related genes, fast-evolving sites/windows only fall within IDRs, but not other structured domains (**Figure 3**, right column).

We also investigated whether varying IDR properties of heterochromatin-related genes contribute to the evolutionary differences between gene categories by estimating the percentage of amino acids falling within predicted IDR domains (% of IDRs). A higher % of IDRs has been found in proteins involved in the formation of phase-mediated cellular domains (e.g., membrane-less organelles (Sawyer et al. 2019) and the heterochromatin domain (Guthmann et al. 2019)). We found that the % of IDR for heterochromatin-related genes in *D. melanogaster* is significantly greater than that of random genes (*Mann-Whitney U test, p* = 0.00214), which is especially true for *Su(var*) and heterochromatin proteins (*Mann-Whitney U test, p* = 0.0448 (*Su(var)s*) and 0.0141 (heterochromatin proteins); **SFigure 4A**). There is a lack of difference in % of IDR between our candidate genes and Polycomb control (*Mann-Whitney U test, p >* 0.05 for all tests, **SFigure 4A**), which may be due to the fact that Polycomb proteins were shown to also undergo phase separation (Tatavosian et al. 2019). Surprisingly, despite previous reports on the reduced evolutionary constraints in IDRs (Brown et al. 2002; Khan et al. 2015), the % of IDRs does not differ between heterochromatin-related genes with or without evidence of positive selection (*Mann-Whitney U test, p > 0.05*; **SFigure 4B**).

**Figure 4.**
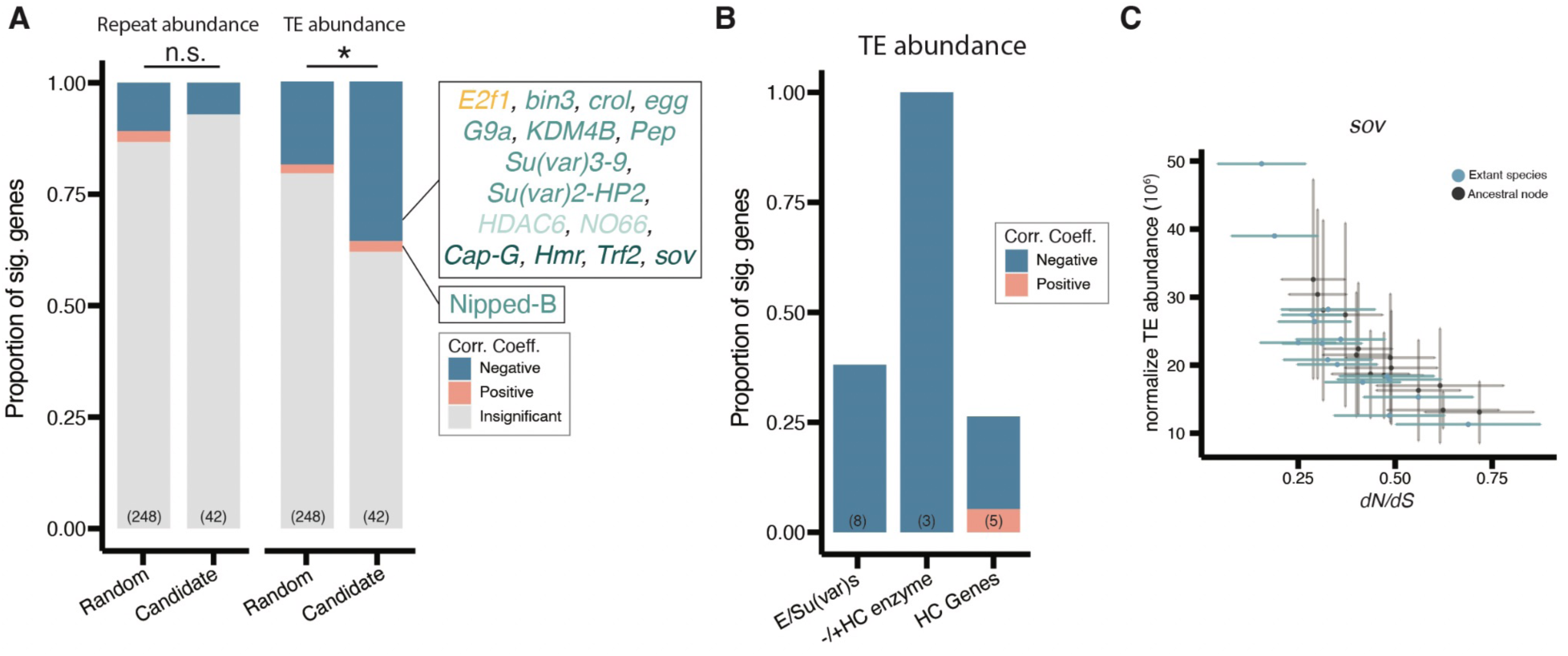
Extensive negative associations between *dN/dS* of heterochromatin-related genes and genomic TE abundance. (A) Stacked bar plots showing the proportion of positively selected genes whose *dN/dS* correlates with repeat abundance (left) or TE abundance (right) for heterochromatin-related genes and randomly sampled genes. The color of the genes in the side box represents the gene’s category: *E(var)s* (yellow), *Su(var)s* (green), histone-modifying enzyme enhancing heterochromatin (light green), and heterochromatin protein (dark green). (B) Bar plots showing the proportion of positively selected heterochromatin-related genes whose *dN/dS* significantly coevolve with TE abundance for different gene categories. (C) X-Y plots showing the associations between *dN/dS* (X-axis) and normalized TE abundance for an example gene (*sov*). Blue points are estimates for the extant species, and gray points are for internal nodes. Lines across each dot denote 95% confidence intervals for *dN/dS* (extant species and internal nodes) and normalized TE content (internal nodes). *Binomial test: n.s. p >* 0.05, \**p* < 0.05.

To further investigate the possible associations between % of IDRs and the evolution of candidate genes, we estimated the phylogenetic signals (Blomberg and Garland Jr 2002) of % of IDRs across studied species. A small *Blomberg’s K* (Blomberg et al. 2003), our chosen index for phylogenetic signal, suggests % of IDR evolves faster than that of random expectation (Brownian motion of trait evolution; (Kamilar and Cooper 2013)). Intriguingly, we found no difference in *K* values between heterochromatin-related genes and control genes, among categories of heterochromatin-related genes that showed different rates of adaptive evolution, and between heterochromatin-related genes with and without evidence of positive selection over long evolutionary time scales (**SFigure 4C and 4D**; *Mann-Whitney U test, p >* 0.05 for all comparisons). These observations indicate that, despite the observed pervasive adaptive evolution of heterochromatin-related genes, their % of IDRs evolves similarly to that of other genes in the genome. In summary, positive selection acts on both structured domains with diverse functions as well as IDRs of heterochromatin-related genes, while stabilizing selection might have maintained the IDR content of these genes in the face of rapidly changing amino acid sequences.

### Rates of protein evolution of heterochromatin-related genes significantly associate with genomic TE abundance

The antagonistic interaction with repetitive sequences enriched in heterochromatin was suggested as the driver for the rapid evolution of a handful of heterochromatin-related genes (Vermaak et al. 2005; Satyaki et al. 2014; Helleu and Levine 2018). To test whether this conjecture may be broadly applicable to heterochromatin-related genes, we estimated the correlation between the rates of amino acid substitution (*dN/dS*) of heterochromatin-related genes and the changes in abundance of heterochromatic repetitive sequences using methods developed in (Lartillot and Poujol 2011). This approach corrects for the phylogenetic nonindependence of the quantitative traits of interest (here, repeat abundance) and models their evolution following the Brownian process. To estimate repeat abundance, we performed Illumina sequencing with PCR-free library preparation to avoid the sequencing bias against AT-rich sequences, which is commonly found in repetitive sequences (Wei et al. 2018). Using these sequencing data, we quantified the genomic abundance of satellite repeats (both simple and complex satellites) and TEs, both which should be dominated by the DNA content of heterochromatin (see Materials and Methods).

Among the 42 heterochromatin-related genes with evidence of positive selection over the long evolutionary time scale, nearly forty percent (38.1%) of them have rates of protein evolution tracking changes in TE abundance across species, with significant (*p* < 0.05) phylogenetically controlled correlations between *dN/dS* and TE abundance (**Table S3**). This proportion is much higher than that of the 248 random genes with accelerated rates of protein evolution (20.5%, *Fisher’s Exact test*, *p* = 0.0173, *Binomial test*, *p* = 0.0112, **Figure 4A**).

Intriguingly, 93.8% of these correlations of the heterochromatin-related genes are in the negative direction, which indicates that these genes evolved faster in species with lower TE abundance (see **Figure 4C** for an example). We also examined this proportion separately for each category of heterochromatin-related genes (**Figure 4B**). Quite strikingly, even though only a small number of histone-modifying enzymes were found to have evidence of positive selection (26.1%), *all* of their rates of protein evolution were significantly associated with TE abundance (**Figure 4B**). These histone-modifying enzymes include all three H3K9me2/3 methyltransferase (*egg, G9a*, and *Su(var)3-9*), *NO66* (H3K4me3 demethylase), *KDM4B* (H3K9me2/3 demethylase), and *HDAC6* (zinc-dependent deacetylase) (**Table S3**). In stark contrast to analysis focusing on TE abundance, only 7.14% of heterochromatin-related genes with evidence of positive selection show a significant correlation between *dN/dS* and the abundance of total satellite repeats, a proportion that is not significantly different from that of randomly sampled genes (**Figure 4A**, *Fisher’s Exact Test p =* 0.322 and *Binomial test p =* 0.360).

Analysis focusing on the abundance of only simple satellites gave consistent results (**Figure S5,** *p* > 0.05 for both *Fisher’s Exact Test* and *Binomial test*). Overall, we found that the rates of protein evolution of heterochromatin-related genes negatively correlated with the abundance of TEs, but not total repeats, across species, suggesting a possible source of selective force shaping the evolution of these genes (see Discussions).

## Discussion

Heterochromatin is a highly conserved cellular compartment with essential functions across complex eukaryotes (Allshire and Madhani 2018; Janssen et al. 2018). Nevertheless, our evolutionary analyses revealed that genes involved in heterochromatin function are highly labile, exhibiting pervasive evidence of rapid evolution both in the forms of amino acid substitutions and gene copy number changes at both short and long evolutionary time scales. Importantly, the rapid evolution of these genes is likely driven by functions specific to constitutive heterochromatin, instead of mechanisms general to proteins interacting with repressive chromatin. Evidence of positive selection on protein evolution is especially prominent for heterochromatin-related genes that should enhance heterochromatin function and, strikingly, *all* three methyltransferases responsible for the enrichment of H3K9me2/3, the characteristic histone modification of heterochromatin, are under positive selection. Our further characterization of the various aspects of the evolution of heterochromatin-related genes provided an important avenue to identify the possible source of selective forces acting on heterochromatin, which we discussed below.

Close examinations of the signatures of positive selection of heterochromatin-related genes found them not only located within structured domains with well-known functions, but also inside unstructured IDRs (**Figure 3**). IDRs were recently found to be critical for the phase properties and thus functions of heterochromatin (Larson et al. 2017; Strom et al. 2017), and our observation suggests that positive selection could also act on heterochromatin functions mediated by such properties. Interestingly, despite the previous suggestions that IDRs are evolutionarily less constrained (Brown et al. 2002; Khan et al. 2015), the proportion of IDR sequences in positively selected heterochromatin-related genes is similar to other candidate genes (**Figure 4B**). Even more, we identified strong phylogenetic signals that are consistent with the presence of stabilizing selection preserving the % of IDRs even for heterochromatin-related genes under strong positive selection. Such findings may indicate that the % of IDRs of a protein, but not the underlying amino acid sequence, is evolutionarily constrained and plays an essential role in the function of studied heterochromatin-related genes. Indeed, a few IDRs with rapidly evolving sequences were found to possess conserved molecular features (Moesa et al. 2012; Zarin et al. 2019), with some of them experimentally demonstrated to be functionally equivalent (Zarin et al. 2017). This raises another question of *why* the underlying amino acid sequences rapidly evolve, considering the need to maintain the IDR content of heterochromatin-related genes (see below).

Several of our findings suggest that TEs, but not satellite repeats, must be involved in the pervasive rapid evolution of heterochromatin-related genes. First, the associations between rates of protein evolution of heterochromatin-related genes and repeat abundance were mainly observed for TEs. In addition, many heterochromatin-related genes under positive selection have functions related to TE suppression. In particular, the interactions between the protein products of several of these genes (e.g., *del, moon, rhi, Trf2*, and *sov* (Klattenhoff et al. 2009; Mohn et al. 2014; Andersen et al. 2017; Andreev et al. 2022)) and repressive H3K9me2/3 are responsible for licensing piRNA clusters, which are TE-enriched loci generating the majorities of piRNAs targeting TEs and located within pericentromeric heterochromatin. Similar to the rapid evolution of the DNA and protein components of centromeres (i.e., the centromeric drive hypothesis, (Henikoff et al. 2001; Malik et al. 2002)), changes in heterochromatic TE sequences may alter their interactions with proteins encoded by heterochromatin-related genes.

Consequently, selection may favor evolutionary changes in proteins that revert the strength of DNA-protein interaction to that prior to the alteration of TE sequences. Subsequent changes in TE sequences could initiate another cycle of this interaction and ultimately drive the fast evolution of genes involved. However, only some of the identified signatures of positive selection fall within nucleic acid binding domains (**Figure 3**), and the majority of the positively selected heterochromatin-related genes lack sequence specificity. Moreover, unlike viruses, the “success” of TEs is tightly intertwined with host fitness, owing to the fact that they propagate in the host germline and are inherited vertically (Haig 2016). Accordingly, an arms race between TEs and host proteins suppressing them was suggested to be unlikely to drive the pervasive adaptive evolution of host genes (Blumenstiel et al. 2016; Cosby et al. 2019).

What might have driven the fast evolution of heterochromatin-related genes then? The genomic autoimmunity hypothesis (Blumenstiel et al. 2016) provides a plausible explanation—molecular mechanisms suppressing TE activities are expected to be constantly juggling between maximizing TE silencing while minimizing inadvertent off-target suppression of functional genes, and this alternating selective pressure could drive the rapid evolution of genes involved. Under this model, for example, variants of heterochromatin-related genes that enhance the generation of piRNAs from piRNA clusters, and thus strengthen TE silencing, may lead to the inadvertent production of piRNAs that target other functional elements, a possibility with empirical support (Andersen et al. 2017; Kelleher 2021). Selection will then instead favor variants increasing the stringency of this licensing to avoid fitness costs of the off-target effect. The repeated alternation of selective targets could accordingly drive the rapid evolution of heterochromatin-related genes.

The genomic autoimmunity hypothesis is even more suitable to explain the rapid evolution of heterochromatin-related genes involved in modulating the intrinsic and unique molecular properties of heterochromatin—a tendency to “spread” to nearby loci in a sequence-independent manner. The positive feedback between writers and readers of H3K9me2/3 promotes the propagation of repressive chromatin marks (Allshire and Madhani 2018; Bell et al. 2023), and the extent of this spreading depends on the concentrations of readers and writers at the suppressed loci (Locke et al. 1988). Heterochromatin-mediated silencing thus needs to be carefully balanced to prevent inadvertent silencing of functional elements while maintaining sufficient suppression at the euchromatin-heterochromatin boundaries. Interestingly, such a balance needs to be maintained not only at the ends of chromosomes (around pericentromeric and subtelomeric heterochromatin), but also around epigenetically silenced, H3K9me2/3-enriched TEs in the euchromatic genome. The spreading of repressive marks from silenced euchromatic TEs into functional genes is widely documented (reviewed in (Choi and Lee 2020)) and has been inferred to impair individual fitness (Lee 2015; Lee and Karpen 2017; Huang et al. 2022). Consistently, several genes known to directly mediate TE epigenetic silencing are found to be under positive selection in our analysis (e.g., *wde, Su(var)2-10,* (Ninova et al. 2020)). In addition, histone-modifying enzymes, which are directly involved in the reader-writer positive feedback loops, should frequently be caught in cycles of alternating selection for enhanced or weakened TE epigenetic silencing. Indeed, we found that *all* three writers for H3K9me2/3 show evidence of rapid evolution (*Su(var)3-9, egg, G9a*) and *all* histone-modifying enzymes with accelerated rates of protein evolution have evidence of coevolving with genomic TE abundance (**Figure 4**). Furthermore, the deleterious off-target effects of TE silencing could result from the long-range spatial interactions between H3K9me2/3-enriched euchromatic TEs and pericentromeric heterochromatin, which is mediated by phase separation mechanisms and is selected against (Y.C.G. Lee et al. 2020). Repeated alteration in selective pressure to ensure these proteins confer sufficient phase properties for proper heterochromatin functions while avoiding such off-target effects may similarly drive rapid changes in protein sequences while preserving the % of IDR for proteins encoded by heterochromatin-related genes, as we have observed.

Intriguingly, we found predominantly negative associations between TE abundance and rates of protein evolution of heterochromatin-related genes, which may initially seem counterintuitive. However, this pattern can be explained by our proposed model that augments the genomic autoimmunity hypothesis by incorporating how the alternating selective pressure on heterochromatin-mediated silencing may concurrently influence genomic TE abundance (**Figure 5A**). Strong heterochromatin-mediated silencing should lead to rampant off-target effects, which not only select *for* variants that weaken the strength of silencing (Huang and Lee 2024), but also select *against* individual TE copies due to the deleterious spreading of repressive marks to TE-adjacent functional sequences. TE abundance should accordingly decrease (**Figure 5A-a**, (Huang et al. 2022)). When variants that reduce heterochromatin-mediated silencing become fixed, the maintenance of heterochromatin at epigenetically silenced TEs weakens, resulting in increased TE replication and more new TE insertions (**Figure 5A-b**). The consequentially increased TE abundance should then drive selection *for* enhanced heterochromatin-mediated silencing, returning to the initial state of strong silencing (**Figure 5B**). Accordingly, as heterochromatin-related genes experience cycles of alternating selection targets and gaining amino acid substitutions, genomic TE abundance also fluctuates. Yet, whether TE abundance eventually increases or decreases depends on the relative strength of selection against TEs with off-target effects and the changes in TE replication rates (**Figure 5B**). Selection coefficients for the harmful off-target effects of TE epigenetic silencing are yet to be estimated, but they could be strong if such effects perturb the expression of nearby vital genes (e.g., (Coronado-Zamora and González 2023)) or disrupt global 3D genome organization (e.g., (Y.C.G. Lee et al. 2020)). On the other hand, replication rates of *Drosophila* TEs are generally low (10^-5^∼10^-4^; (Charlesworth and Langley 1989; Adrion et al. 2017; Wang et al. 2023)). Changes in these rates are likely weaker than selection removing TEs through their off-target effects, leading to decreased TE abundance over cycles of alternating selective pressure on the strength of heterochromatin-mediated silencing and thus giving rise to negative associations between genomic TE abundance and rates of protein evolution between species. It is worth noting that many other processes, such as recent demographic changes (e.g., (Mérel et al. 2021)), also contribute to between-species differences in TE abundance. If these forces systematically influence genomic TE abundance in the species studied (e.g., correlate with the evolution of heterochromatin-related genes), similar associations could arise in the absence of the proposed model.

**Figure 5.**
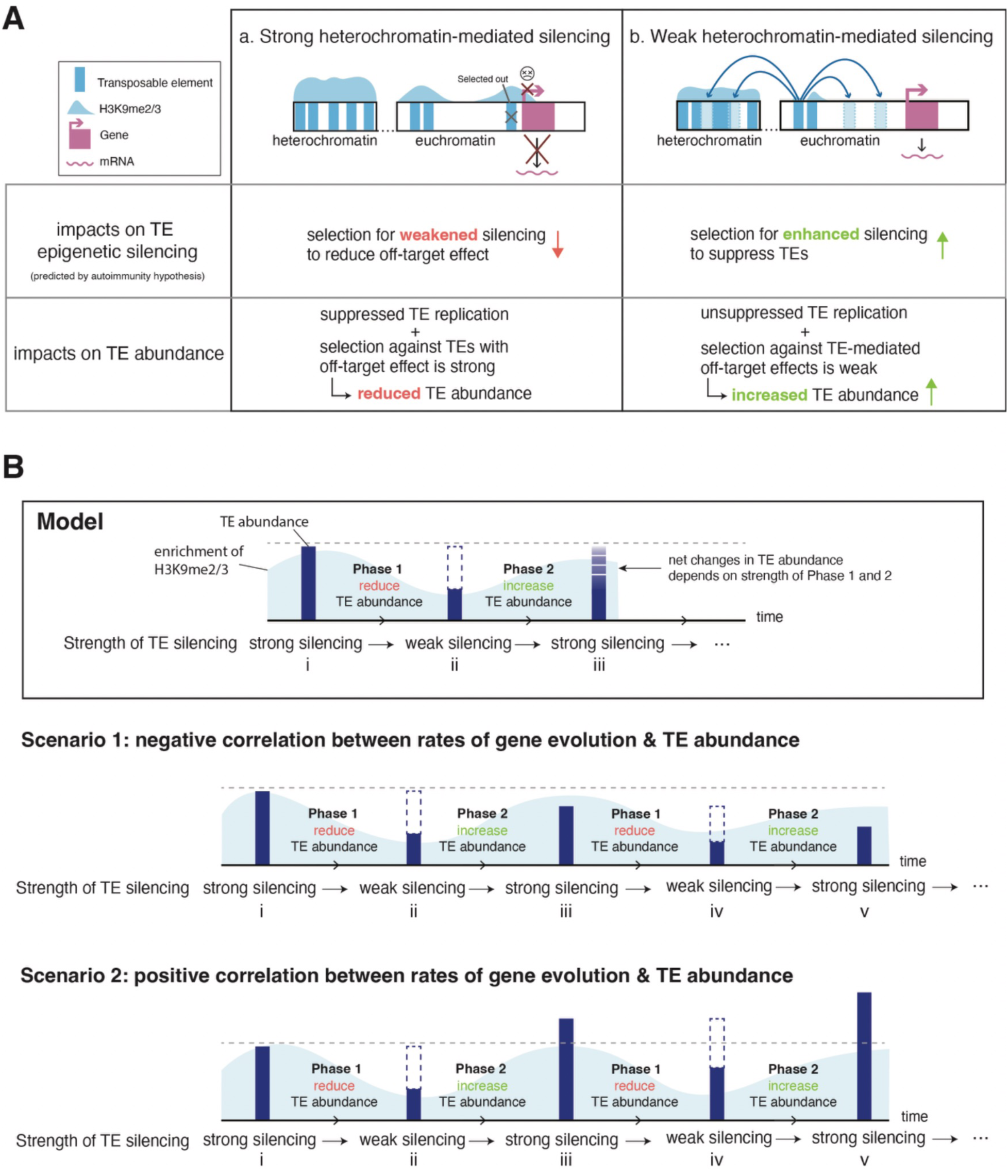
Model for the correlated evolution of heterochromatin-related genes and genomic TE abundance. (A) The predicted impacts of different strengths of heterochromatin-mediated silencing on TE epigenetic silencing and genomic TE abundance. **(A-a)** Strong heterochromatin-mediated silencing results in strong off-target effects due to the spreading of repressive marks from silenced euchromatic TEs, leading to positive selection for variants of heterochromatin-related genes that weaken the strength of silencing. In addition, because of the low rates of TE replication from strong silencing and negative selection against TEs with off-target effects, TE abundance should reduce. (**A-b**) On the other hand, when heterochromatin-mediated silencing is weak, selection against TE-mediated off-target effects is weak and rates of TE replication should be high, leading to increased TE abundance. At the same time, selection may favor variants that enhance TE silencing to reduce rates of TE replication. (**B**) Proposed sequences of events that lead to correlated evolution of heterochromatin-related genes and genomic TE abundance. The model should be applicable to any of the states and the following description starts with strong heterochromatin-mediated silencing (i). TE abundance is expected to reduce due to the suppressed rates of TE replication and selection against TEs with off-target effects (Phase 1). Concurrently, there should be selection for variants of heterochromatin-related genes that weaken heterochromatin-mediated silencing to reduce deleterious off-target effects. When variants that weaken silencing are fixed, the maintenance of TE epigenetic silencing decreases (ii), leading to both reduced occurrence of off-target effects and increased rates of TE replication. TE abundance should thus increase (Phase 2). The high TE abundance and replicative activity would select for variants of heterochromatin-related genes that enhance TE silencing, going back to the initial state of strong heterochromatin-mediated silencing (e.g., iii here). The relative strength of a decrease in TE abundance through selection against off-target effects and increase in TE abundance through TE replication determines whether there are negative (scenario 1) or positive (scenario 2) associations between TE abundance and the evolution of heterochromatin-related genes across species.

Our proposed model for how the need of maintaining a balanced TE silencing drives the fast evolution of heterochromatin-related genes might help explain some intriguing patterns we observed. Majorities of our identified duplicates of heterochromatin-related genes (48.78%) have male-biased expression (**Figure 2**), indicating their potential functional significance in male germlines. While the resolution of sexual genetic conflict (Gallach and Betrán 2011; Wyman et al. 2012), a common explanation for male-biased expression of newly duplicated genes, may underlie such an observation, it may also stem from the need to properly silence the male-specific TE landscape (Chen et al. 2021), resulting in sexually dimorphic chromatin environment. Another intriguing observation is the limited evidence of positive selection for heterochromatin-antagonizing genes (**Figure 1**), which is in stark contrast to the pervasive signatures of adaptive evolution of genes mediating heterochromatin functions. Protein products of heterochromatin-antagonizing genes are not only found at heterochromatin-euchromatin boundaries, but broadly distributed throughout the euchromatic genome (Dimova et al. 2003; Kellner et al. 2012), playing diverse roles in transcriptional regulation (Meignin and Davis 2008; Regnard et al. 2011; Crona et al. 2013). The pleiotropic functional roles of heterochromatin-antagonizing proteins, in both euchromatin and heterochromatin, should constrain their evolution, making them less likely to be caught up in alternating selection for the sensitivity or the specificity of TE silencing. Nevertheless, the small sample size of heterochromatin-antagonizing genes (only 12 genes) could limit our ability to detect positive selection among these genes.

Our observed frequent positive selection acting on heterochromatin-related genes points out the high lability of the molecular feature of this functionally conserved genomic compartment. Instead of the usually assumed “arms race” with the changing repeatome, we proposed that the selective pressure of heterochromatin-related genes might mainly come from a need to maintain a delicate balance of its unique ability to “suppress” and to “spread,” which also consequently influence the evolutionary dynamics of TEs. Signatures of positive selection identified here could serve as an entry point to further investigate how the delicate balance of heterochromatin-mediated silencing may be conferred by vastly changing components between species (e.g., (Rosin and Mellone 2016; Parhad et al. 2017)), providing a fruitful opportunity to further dissect the molecular mechanisms shaping heterochromatin functions and evolution.

## Materials and Methods

### Evolutionary analyses of protein sequences

Coding sequences and genome annotations for 16 studied species (**Figure 1B**) were downloaded from NCBI, with GenBank ID listed in **Table S4**. Because we compiled the list of candidate genes based on *D. melanogaster* literature, we used orthologous information from OrthoDB (Kuznetsov et al. 2023)(last accessed 12/10/2022) to retrieve one-to-one orthologs for *D. melanogaster* candidate genes. For genes without one-to-one ortholog according to OrthoDB, we performed *BLAST* search (see below). The retrieved coding sequences were translated to amino acid sequences, aligned using Clustal Omega (version 1.2.4; (Sievers et al. 2011)), and converted back to codon alignments using PAL2NAL (version 14; (Suyama et al. 2006)).

We performed unpolarized McDonald-Kreitman (MK) tests (McDonald and Kreitman 1991) using polymorphism within a *D. melanogaster* Zambian population (197 strains, (Lack et al. 2015)) and divergence between *D. melanogaster* and *D. simulans.* Following the recommendation of (Lack et al. 2015), we treated genomic regions with non-African ancestry or identity-by-descent as missing data, and only included sites with at least 100 non-missing alleles. To count the number of nonsynonymous and synonymous changes, we used the mutational paths that minimize the number of amino acid changes. Codons with more than two alleles, considered both within species polymorphism and between species divergence, were excluded. Genes with fewer than 100 codons were excluded from the analysis due to insufficient statistical power. A gene is deemed under positive selection if the MK test, whose significance was assessed using *Fisher’s Exact* test, rejected the null hypothesis and the ratio of nonsynonymous to synonymous changes is greater for between-species substitutions than for within-species polymorphism. Sliding window MK tests were performed with windows of 100 codons and 10-codon steps.

We conducted phylogenetic analysis by maximum likelihood (PAML) (Yang 2007) using 16 species to identify candidate genes experiencing positive selection over a long evolutionary time scale. We compared two site models, M8a (*dN/dS* ratio is beta-distributed and not greater than one) and M8 (*dN/dS* > 1), and determined the significance using likelihood ratio tests. The species tree reported in (Suvorov et al. 2022) was used. Sites with > 0.95 BEB posterior probability of coming from the site class with *dN/dS* > 1 (Yang et al. 2005) are considered under recurrent adaptive evolution.

### Evolutionary analysis for gene copy number

To identify genes with varying gene copy numbers, we first used a genome-wide, high throughput search with liberal parameters to identify many potential candidates, followed by careful manual curations. We first used tblastn and reciprocal blastx (Camacho et al. 2009) to search for homologs and paralogs of candidate genes in studied species using *D. melanogaster* amino acid sequence as queries, with the following parameter: e-value < 10^-2^, amino acid identity > 20%, and blast score > 10. We required the best reciprocal blastx hit to be the original *D. melanogaster* query. Each potential CNV was manually validated using reciprocal best blast with more stringent criteria (e-value < 10^-5^), and orthologs and paralogs were distinguished using synteny of flanking genes (**Figure S1**). We further examined the expression levels of candidate CNVs using RNA-seq exon coverage tracks of NCBI Data Viewer and removed those with no expression. Several duplicates identified in *D. suzukii* were filtered due to redundant contigs. For gene loss, we followed the procedures detailed in (King et al. 2019) to confirm the true absence of a gene.

To examine the sex-biased expression, we deemed a gene male-biased if the log2 fold change of the ratio of male and female expression (TPM or FPKM) is >1, female-biased if such value is < –1, and otherwise unbiased. For *D. melanogaster,* we used Insect Genome database (http://www.insect-genome.com/Sexdb/, last retrieved November 2023) and FlyAtlas2 RNA-seq data (Krause et al. 2022) for whole adult males and whole adult females. For the other 15 species, we mapped publicly available transcriptome datasets (**Table S5**) to publicly available genome assemblies with gene annotations using HiSAT2 (v2.2.1 with parameters –exon and – ss to specify the exon positions and splice sites; RRID:SCR_015530; (Kim et al. 2019)), and estimated the expression levels usingStringtie (v2.1.4 with parameters –dta –G to specify annotation files; RRID:SCR_016323; (Kovaka et al. 2019)). Candidate duplicates identified by manual curation but have no annotation or show no expression were excluded from the analysis.

### Domain annotations

We used UniProt (The UniProt Consortium 2023) to annotate known structured domains within the *D. melanogaster* allele of heterochromatin-related genes. We used flDPnn (Hu et al. 2021), which performed superior in the previous benchmark study (Necci et al. 2021), and the predicted binary index for IDRs (disorder propensity cutoff = 0.3) to annotate IDRs for all studied species. To detect the phylogenetic signal of the % of IDR, we computed *Blomberg’s K* statistic using the phylosig function from phytools R package (Blomberg et al. 2003; Revell 2012). The tree structure and branch lengths were obtained from (Suvorov et al. 2022) and generated by treeio R package (Wang et al. 2020).

### Analyses of repetitive sequences and their coevolution with candidate genes

DNA for each studied species was extracted from 40 females using DNeasy Blood & Tissue Kit (Qiagen), following the manufacturer’s protocol. To avoid PCR amplification bias during the preparation of sequencing libraries, which was found to skew the quantification of repeats (Wei et al. 2018), extracted DNA was prepared into Illumina sequencing library with PCR-free protocol and sequenced with 150bp paired-end reads by Novogene (Sacramento, CA).

We used Satellite Repeat Finder (Zhang et al. 2023) to estimate the abundance of total satellite repeat in the Illumina short-read sequencing data. Following the suggested procedures, we first counted K-mers (K=21) in each sample using KMC (ver. 3.2.4; (Kokot et al. 2017)). Contigs of satellites were then generated and mapped back to the source Illumina sequences to estimate the abundance (in bases) of each satellite using minimap (ver. 2.24; (Li 2018)) and Satellite Repeat Finder. We also used K-seek (Wei et al. 2014) to estimate the abundance of simple repeats and reached similar conclusions (see Results).

TE abundance was estimated as the total TE reads from the Illumina sequencing data, with the assumption that TEs in the heterochromatin mainly originated from jumping events of euchromatic TEs and TE abundances in the heterochromatin and euchromatin are highly correlated. To minimize TE annotation bias across species, repetitive sequences from each of the 16 genomes (**Table S4**) were identified using RepeatMasker (version 4.1.0; http://www.repeatmasker.or/) with the Dfam database (Storer et al. 2021), using the command “RepeatMasker –q [genome sequence file] –species drosophila –e hmmer”. TE sequences annotated as LTR, LINE, DNA element, and Unknown categories were obtained from all 16 genomes to create a master TE library. Illumina reads from each species were then mapped to the library with bwa-mem (version 0.7.17; (Li and Durbin 2009)) and viewed by samtools (version 1.15.1, (Li 2011)). The total number of reads mapped to the library, regardless of mapping quality, was considered the number of TE reads. It is worth noting that very few TEs were classified as Unknown category, and inclusion/exclusion of such TEs did not change the results. To compare the abundance of satellite repeats and TEs across samples/species, these estimates were normalized. Illumina reads from each species were mapped to its repeat-masked genome using bwa-mem, with sites with mapping quality lower than 30 filtered (using samtools –q 30). The median read depth for the unmasked regions for each sample was then used to normalize the number of bases for satellite repeats and TEs.

We used Coevolve (Lartillot and Poujol 2011) to estimate the correlation between *dN/dS* and repeat abundance (satellite repeats or TEs) given the tree structure. For each gene, the analysis was performed in duplicate to assess convergence (relative difference < 0.1) and a burn-in of 300 with at least 3,000 MCMC chains to get the final estimated correlation between *dN/dS* and repeat abundance. Because the number of Polycomb genes with significant PMAL tests is small, we only compared the coevolution results of candidate genes to those of 248 randomly chosen genes with accelerated rates of amino acid evolution (i.e., significant for PAML analysis).

## Data and script availability

PCR-free Illumina data has been deposited to SRA under the accession number PRJNA1113679. Scripts used in this study can be found at https://github.com/YuhengHuang87/HC_gene_evo and https://github.com/hilynano/Homology_detection.

## Supporting information

SFigures

STables

## Acknowledgments

We thank the High Performance Cluster at UC Irvine for computational resources and Harsh Shukla for helping with the flDPnn software. We appreciate Aniek Janssen, Andrea Betancourt, and Adriana Ludwig for their helpful comments on the manuscript. CHC was supported by Damon-Runyon Cancer Research Foundation postdoctoral fellowship DRG 2438-21 and NIH R01-GM74108 (to Hamit S Malik), SUC was supported by NIH R35GM139653 (to Gary H. Karpen), YH, LL, JM, and YCGL were supported by NIH R35-GM14292 (to YCGL).

## Supplementary Materials

**Table S1. List of candidate and control genes**

Names, FBgn, categories, and references of candidate heterochromatin-related genes and Polycomb control genes. Results of evolutionary tests of each of the genes are also included.

**Table S2. Duplicates of heterochromatin-related genes identified in this study**

**Table S3. Correlation between repeat abundance and dN/dS ratio for positively selected heterochromatin-related genes**

*\*p < 0.05, **p< 0.01, ***p< 0.001*

**Table S4. NCBI assembly number of reference genomes used in this study**

**Table S5. Transcriptome resources and information used in this study**

**Figure S1. Synteny information for genes with duplicates found in more than one species** The following figures show the NCBI genome browser tracks for genes with duplicates (red boxes) found in more than one species. Genes flanking the duplicates (either dark or light blue boxes) were used to infer the synteny of duplicates between species.

**Figure S2. Evolutionary history of *mof* duplicates**

The generation of *mof* CNVs likely involved a duplication in the common ancestor of the oriental linage (green), followed by a subsequent loss in the subsets of lineages (red) and a lineage-specific duplication event (green leading to *D. kikkawai*), resulting in the paraphyletic presence of a *mof* duplicate. Numbers next to species indicate the number of gene copies; species without labeled numbers have one gene copy.

**Figure S3. Proportion of genes with more than one significant evolutionary tests**

Barplots showing the proportion of genes found to be under positive selection and/or fast evolve with at least one (A) or at least two (B) conducted evolutionary tests. The numbers of genes in each category are shown in parentheses. –HC enzyme: histone-modifying enzymes weakening H3K9me2/3 enrichment; +HC enzyme: histone-modifying enzymes enhancing H3K9me2/3 enrichment. *Binomial test*: *\*\*\*p <* 0.001.

**Figure S4. The percentage of intrinsically disordered regions (% of IDRs) among gene categories**.

(A and B) Violin plots comparing the % IDR of *D. melanogaster* proteins for heterochromatin-related genes and the Polycomb control (A) and for heterochromatin-related genes with and without evidence of positive selection over both long and short evolutionary time scales (B). (C and D) Violin plots comparing *Blomberg’s K* for % of IDR across species for heterochromatin-related genes and the Polycomb control (C) and between heterochromatin-related genes with and without evidence of positive selection over a long evolutionary time scale (D). *Mann-Whitney U test*: \**p <* 0.05 and *n.s. p >* 0.05.

**Figure S5. Associations between *dN/dS* of heterochromatin-related genes and the abundance of simple repeats**.

Stacked bar plots showing the proportion of positively selected genes whose *dN/dS* correlates with the abundance of simple satellite repeats for heterochromatin-related genes and randomly sampled genes. The numbers of genes in each category are in parentheses. *Binomial test: n.s. p >* 0.05.

