## Supplementary material for "Prevalent fast evolution of genes involved in heterochromatin functions": SFigures

**Figure S1. Synteny information for genes with duplicates found in more than one species**

The following figures show the NCBI genome browser tracks for genes with duplicates (red boxes) found in more than one species. Genes flanking the duplicates (either dark or light blue boxes) were used to infer the synteny of duplicates between species.

### JIL-1

We identified two duplicates, with JIL-1-like1 present in *D. kikkawai* and *D. serrata* and JIL-1-like2 present in only *D. serrata*.

#### 1. JIL-1-like1

*D. kikkawai* duplicate: LOC108075992 – red box
*D. serrata* duplicate: LOC110180208 – red box
Neighboring genes in synteny to the duplicate:
LOC108075994 (*D. kikkawai*) and LOC110180209 (*D. serrata*) – dark blue box
LOC108076086 (*D. kikkawai*) and LOC110180206 (*D. serrata*) – light blue box

LOC108075992 (*D. kikkawai* duplicate)

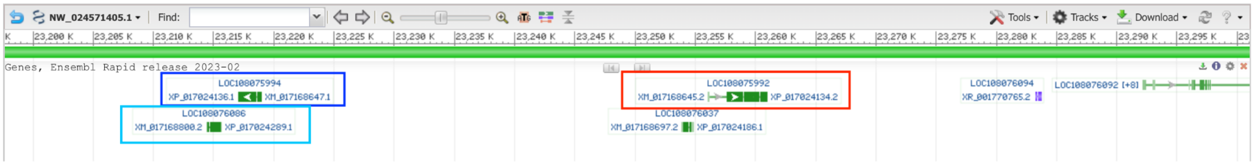


LOC110180208(*D. serrata* duplicate)

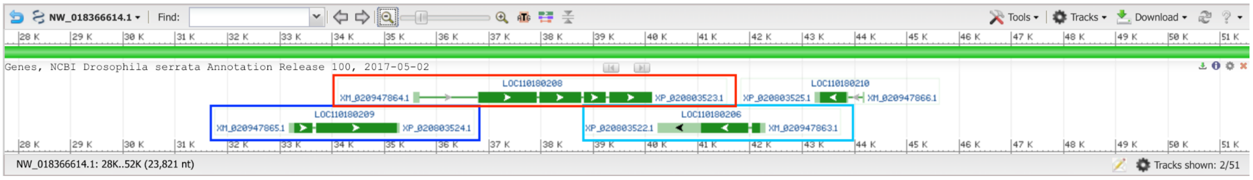


#### 2. JIL1-like2

*D. serrata* duplicate: LOC110182484 – red box
No neighboring genes in synteny to other duplicates

LOC110182484 (*D. serrata* duplicate)

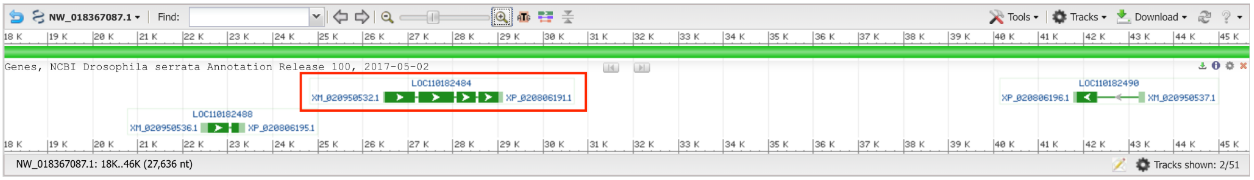

### mof

We identified two duplicates, with mof-like1 present in *D. kikkawai* and mof-like2 present in *D. elegans, D. rhopaloa,* and *D. ficusphila*.

#### 1.mof-like1

*D. kikkawai* duplicate: LOC108075875 – red box

No neighboring genes in synteny to other duplicates

LOC108075875 (*D.kikkawai* duplicate)

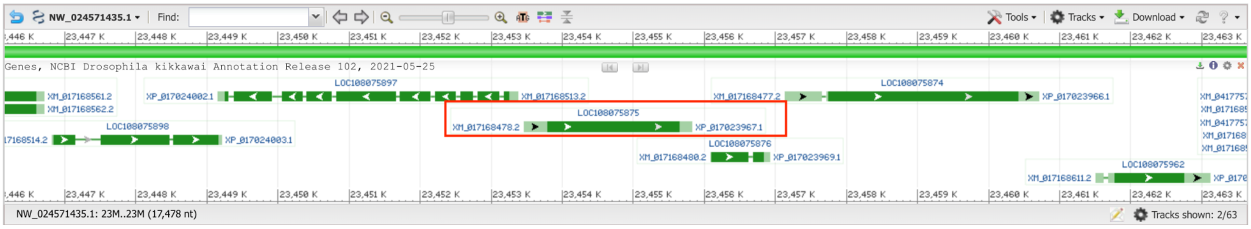


#### 2. mof-like2

*D. elegans* duplicate: LOC108148462 – red box

*D. rhopaloa* duplicate: LOC108046768 – red box

*D. ficusphila* duplicate: LOC108102269 – red box

Neighboring genes in synteny to the duplicate:

LOC108148658 (*D.elegans*), LOC108046767 (*D.rhopaloa*), LOC108102302 (*D.ficusphila*) – dark blue box

LOC108148911 (*D.elegans*), LOC108046809 (*D.rhopaloa*), LOC108102297 (*D.ficusphila*) – light blue box

LOC108148462 (*D.elegans* duplicate)

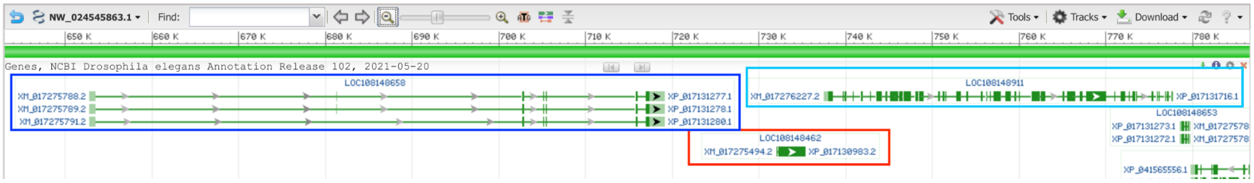


LOC108046768 (*D.rhopoloa* duplicate)

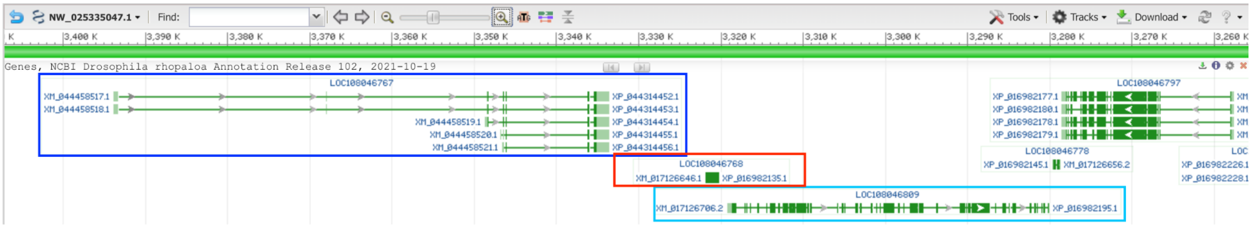


LOC108102269 (*D.ficusphila* duplicate)

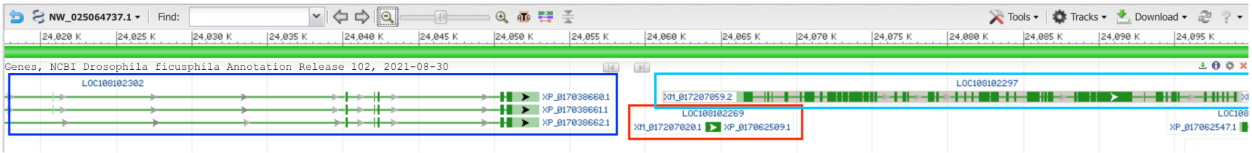

### <u>del</u>

We identified one duplicate, with del-like1 present in *D. simulans* and *D. sechellia*.

#### 1.del-like1

*D. simulans* duplicate: LOC6733043 – red box
*D. sechellia* duplicate: LOC6618245 – red box
Neighboring genes in synteny to the duplicate:
LOC6733041 (*D. simulans*) and LOC6618243 (*D. sechellia*) – dark blue box
LOC6733045 (*D. simulans*) and LOC6618246 (*D. sechellia*) – light blue box

LOC6733043 (*D. simulans* duplicate)

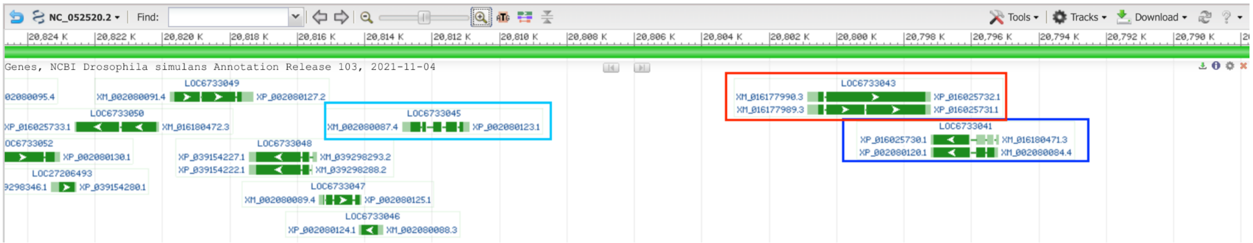


LOC6618245 (*D. sechellia* duplicate)

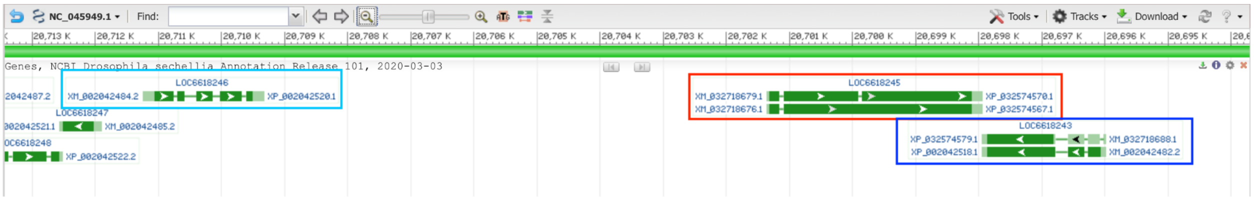

### Nipped-B

We identified two duplicates, with Nipped-B-like1 present in *D. suzukii* and Nipped-B-like2 present in *D. yakuba*.

#### 1.Nipped-B-like1

*D. suzukii* duplicate: LOC108017528 – red box

No neighboring genes in synteny to the other duplicate

LOC108017528 (*D. suzukii* duplicate)

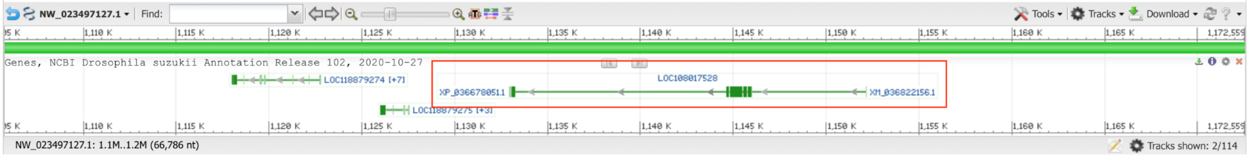


#### 2. Nipped-B-like2

*D. yakuba* duplicate: LOC6539275 – red box

No neighboring genes in synteny to the other duplicate

LOC6539275 (*D. yakuka* duplicate)

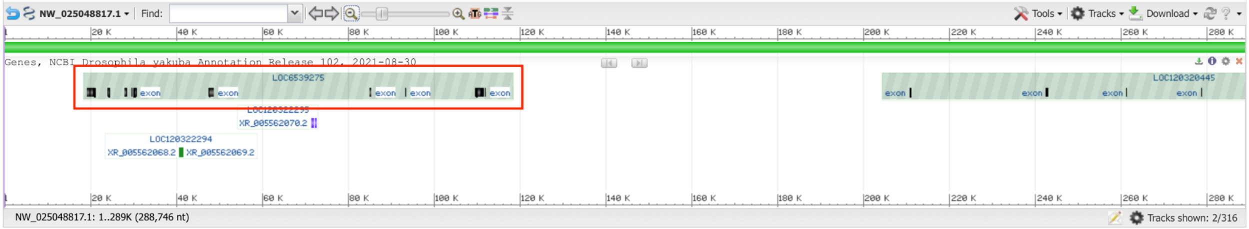

## Pp1-87B

We identified three duplicates, with Pp1-87B-like1 present in *D. kikkiwai* and *D. serrata*, Pp1-87B-like2 present in *D. kikkiwai* only, and Pp1-87B-like3 present in *D. serrata* only.

#### 1.Pp1-87B-like1

*D. kikkawai* duplicate: LOC108080856 – red box
*D. serrata* duplicate: LOC110189278 – red box
Neighboring genes in synteny to the duplicate:
LOC108080855 (*D. kikkwai*) and LOC110189277 (*D. serrata*) – dark blue box
LOC108080857 (*D. kikkwai*) and LOC110189582 (*D. serrata*) – light blue box

LOC108080856 (*D.kikkwai* duplicate)

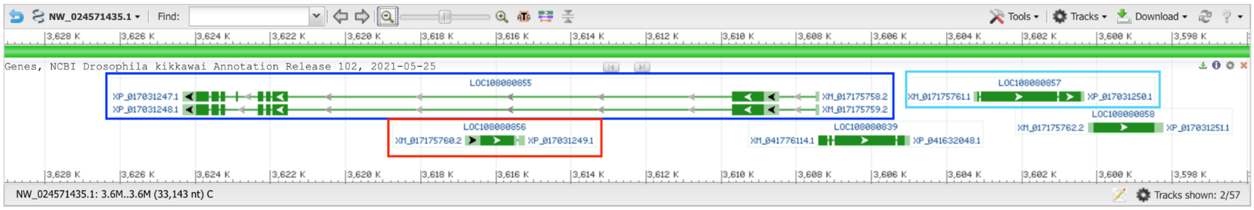


LOC110189278 (*D.serrata* duplciate)

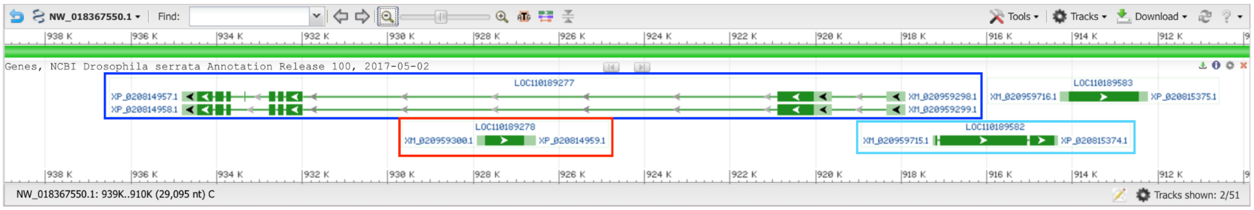


#### 2. Pp1-87B-like2

*D. kikkawai* duplicate: LOC108072267 – red box
No neighboring genes in synteny to the other duplicates

LOC108072267 (*D.kikkawai* duplicate)

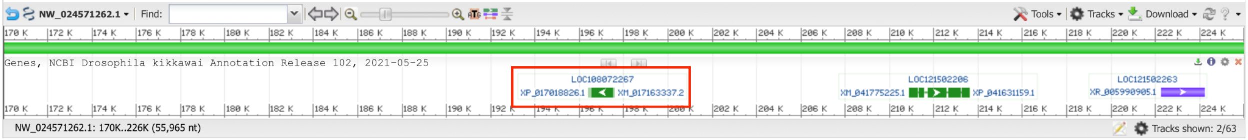


#### 3. Pp1-87B-like3

*D. serrata* duplicate: LOC110178105 – red box
No neighboring genes in synteny to the other duplicates

LOC110178105 (*D. serrata* duplciate)

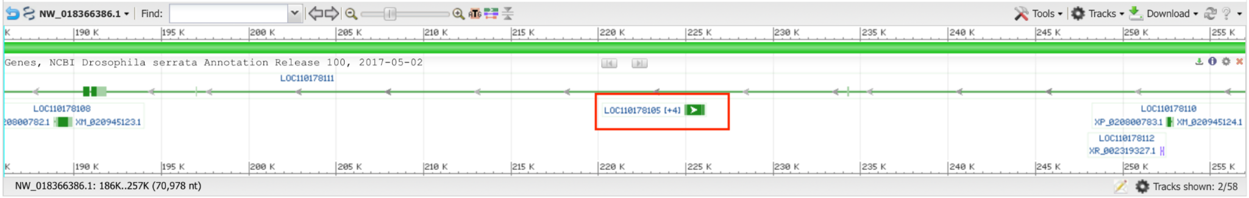

**snf**

We identified one duplicate, with snf-like1 present in *D. takahashii* and *D. eugracillis*.

**1.snf-like1**

*D. takahashii* duplicate: LOC108067435 – red box

*D. eugracilis* duplicate: LOC108104096 – red box

Neighboring genes in synteny to the duplicate:

LOC108067429 (*D. takahashii*) and LOC108104036 (*D. eugracilis*) – dark blue box

LOC108067497 (*D. takahashii*) and LOC108104095 (*D. eugracilis*) – light blue box

LOC108067435 (*D.takahashii* duplicate)

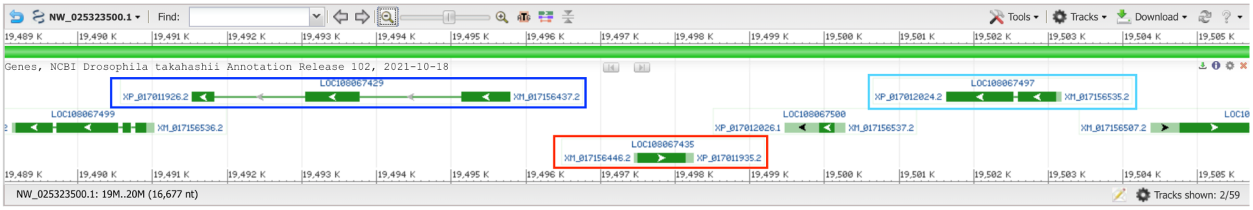


LOC108104096 (*D. eugracilis* duplicate)

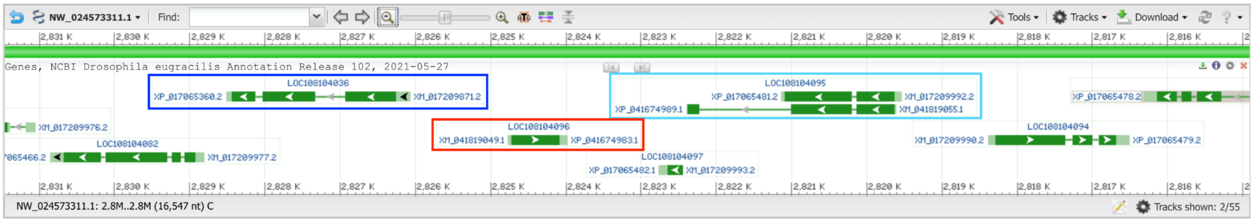

**Figure S2. Evolutionary history of *mof* duplicates**

The generation of *mof* CNVs likely involved a duplication in the common ancestor of the oriental linage (green), followed by a subsequent loss in the subsets of lineages (red) and a lineage-specific duplication event (green leading to *D. kikkawai*), resulting in the paraphyletic presence of a *mof* duplicate. Numbers next to species indicate the number of gene copies; species without labeled numbers have one copy of *mof*.

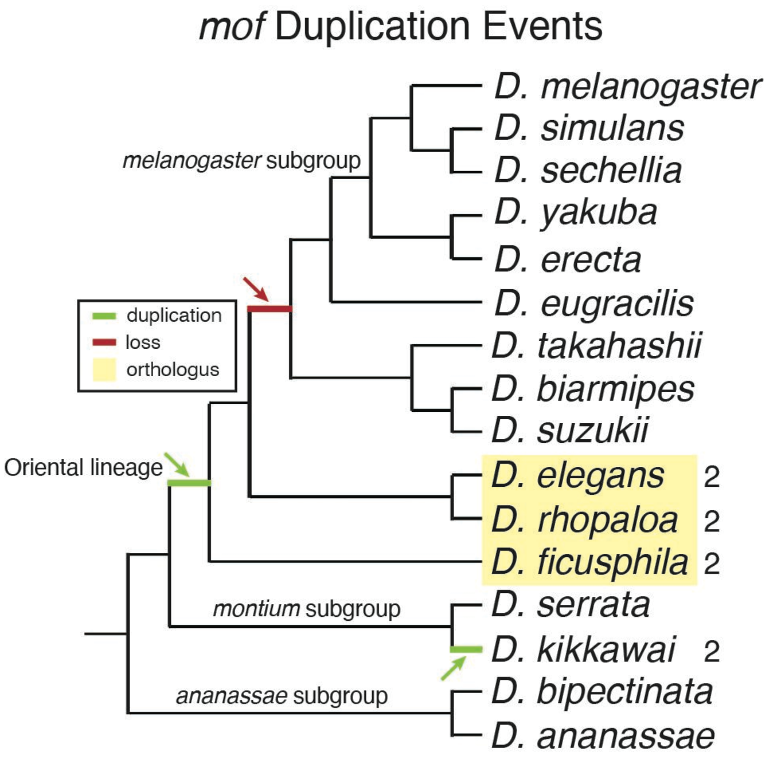

**Figure S3. Proportion of genes with more than one significant evolutionary tests**

Barplots showing the proportion of genes found to be under positive selection and/or fast evolve with at least one (A) or at least two (B) conducted evolutionary tests. The numbers of genes in each category are shown in parentheses. -HC enzyme: histone-modifying enzymes weakening H3K9me2/3 enrichment; +HC enzyme: histone-modifying enzymes enhancing H3K9me2/3 enrichment. *Binomial test*: ***$p$ < 0.001.

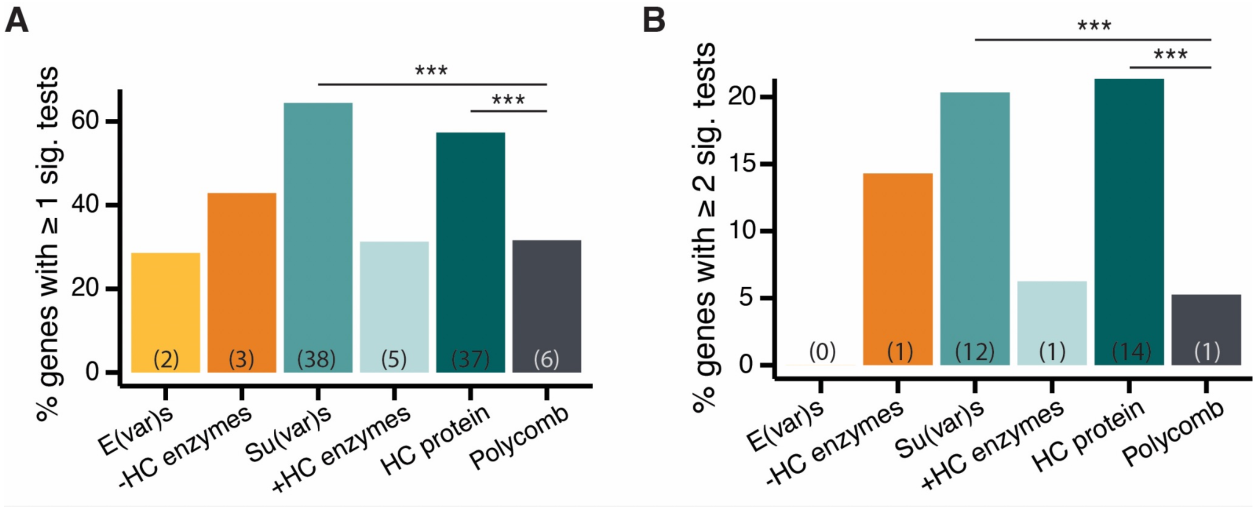

**Figure S4. The percentage of intrinsically disordered regions (% of IDRs) among gene categories.**

(A and B) Violin plots comparing the % IDR of *D. melanogaster* proteins for heterochromatin-related genes and the Polycomb control (A) and for heterochromatin-related genes with and without evidence of positive selection over both long and short evolutionary time scales (B). (C and D) Violin plots comparing *Blomberg's K* for % of IDR across species for heterochromatin-related genes and the Polycomb control (C) and between heterochromatin-related genes with and without evidence of positive selection over a long evolutionary time scale (D). *Mann-Whitney U test*: *$p < 0.05$ and *n.s.* $p > 0.05$.

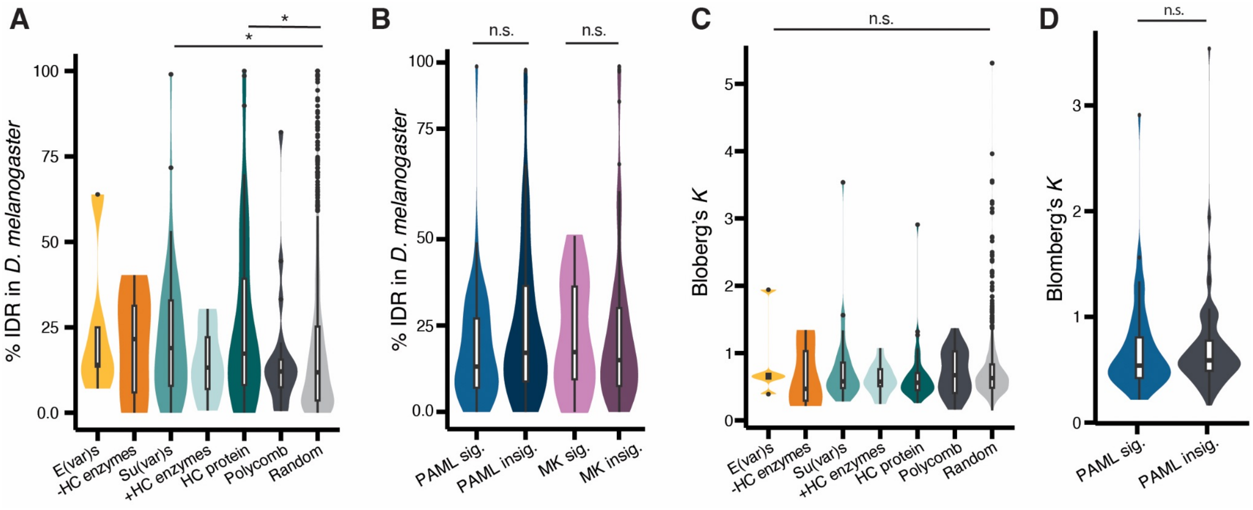

**Figure S5. Associations between *dN/dS* of heterochromatin-related genes and the abundance of simple repeats.**

Stacked bar plots showing the proportion of positively selected genes whose *dN/dS* correlates with the abundance of simple satellite repeats for heterochromatin-related genes and randomly sampled genes. The numbers of genes in each category are in parentheses. *Binomial test: n.s.* $p > 0.05$.

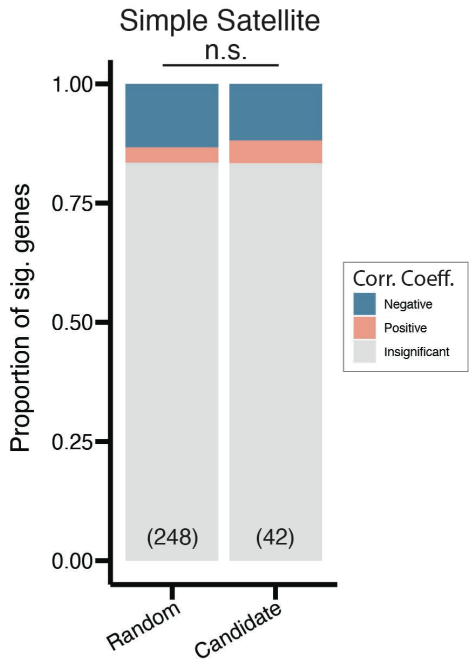
